# NGS-based assay for frequent newborn inherited diseases: from development to implementation

**DOI:** 10.1101/050419

**Authors:** T. Simakova, A. Bragin, M. Zaytseva, C. Clemente, M. Lewicka, J.C. Machado, J.L. Costa, M. Hughes, C. Hertz-Fowler, N. Petrova, A. Polyakov, R. Zinchenko, E. Kondratyeva, A. Pavlov

## Abstract

NGS is a powerful tool for the diagnostics of inherited diseases. A number of studies devoted to the development and validation of targeted NGS panels are published. Here we present not only development and validation of an assay, but report our experience on introduction of a new approach into the real clinical practice. The assay is intended for the diagnostics of frequent newborn inherited diseases: cystic fibrosis, phenylketonuria and galactosemia. The analysis is performed on the Ion PGM™ sequencing platform and allows the detection of single-nucleotide variations as well as copy number variants. We developed the software performing data quality control, providing decision-support variant annotation and generating the medical report that enables clinical application of the assay. Analytical validation of the assay was performed by bi-directional Sanger sequencing of the most part of the targeted region. Clinical validation was performed by multicenter blind testing of clinical and control samples. Sensitivity and specificity of the assay are above 99%. We have developed statements for test ordering, test acquisition form and practical recommendations for the results interpretation. The test has been successfully applied for the confirmatory diagnostics in a clinical laboratory during a year. Thus, the developed assay is a comprehensive ready-to-use CE-IVD solution for clinical diagnostics.

## INTRODUCTION

NGS is a promising tool for the diagnostics of recessive monogenic disorders. Generally, these disorders are conditioned by two pathogenic alleles occurring at the same locus of homologous chromosomes. Thus, identification of two pathogenic alleles may be used for the diagnosis confirmation. Many frequent monogenic disorders are genetically heterogeneous with hundreds known causative variants. For such diseases high-throughput NGS technology is the most appropriate and informative method. High performance of different NGS platforms for clinically significant variant detection has been revealed in a number of recent proof-of-concept studies (**Elliott 2012**, **Umbarger 2013**, **Abou Tayoun 2013**, **Lee 2014**, **Grosu 2014**, **Cao 2014**, **Gu 2014**, and **Trujillano 2015**). Even variants difficult for detection, such as copy number variations (CNV), short tandem repeat (STR) variations and variants in homopolymer regions, can be efficiently detected using targeted sequencing technology. However, the adoption of new technology in a routine clinical practice is limited for the diagnostics of monogenic disorders. This is caused by difficulties in assay validation and results interpretation since unlimited number of variants including rare and unknown may be detected. Laboratories offer NGS-based assays as laboratory-developed tests (LDT) whose performance characteristics are valid only in manufacturing laboratory. Results interpretation is entrusted to a board-certified clinical geneticist. (**Kulkarni 2015**).

This study is aimed at the development and validation of an *in vitro* NGS-based assay acceptable for clinical diagnostics. Therefore, our efforts were focused on quality control, analytical and clinical validation, variant annotation, results interpretation and introduction the assay workflow into the clinical lifecycle. We selected three most frequent monogenic disorders tested within the framework of Russian newborn screening as assay targets: cystic fibrosis (CF), phenylketonuria (PKU) and galactosemia (GAL). We developed assay workflow for the Ion PGM™ sequencing platform and data analysis pipeline using specific software, performed extensive clinical trials and introduced the assay in a clinical laboratory.

## MATERIALS AND METHODS

### Panel design

Ion AmpliSeq Designer v3.4.2 software (Life Technologies, Carlsbad, CA, USA) was used for primer design for targeted regions. Targeted regions were designed to cover all selected significant variants (see **Assay design)**.

### Assay workflow

DNA was isolated from whole blood or saliva using the Quick-gDNA MiniPrep kit (Zymo Research, Irvine, CA, USA). DNA concentrations were determined using a Qubit^®^ 2.0 fluorometer and Qubit^®^ dsDNA HS assay kit (Life Technologies). The minimal amount of DNA required for further analysis was 20 ng (6 μl at a concentration of 3.3 ng/μl).

DNA libraries were constructed with Ion AmpliSeq™ Library Kit v2.0 and two custom 2x Ion AmpliSeq Primer Pools according to the manufacturer’s instructions. PCR-reactions were performed in a MyCycler™ thermal cycler (Bio-Rad, Hercules, CA, USA). Amplicons of the same DNA from the two pools were pooled together and treated with FuPa Reagent (Life Technologies) to partially digest the primers and phosphorylate the amplicons. Ion Xpress™ barcoded adaptors (Life Technologies) were ligated to the amplification products. The resulting DNA libraries were purified using Agencourt AMPure™ XP Reagents (Beckman Coulter, Brea, CA, USA) according to the manufacturer’s instructions. Purified libraries were then quantified using a Qubit^®^ 2.0 fluorometer and Qubit^®^ dsDNA HS assay kit (Life Technologies). The final libraries concentrations were diluted with Low TE buffer (Life Technologies) to 100 pM and mixed. Template preparation was performed using Ion One Touch™ System and Ion OneTouch™ 200 Template Kit v2 DL (Life Technologies) according to the manufacturer’s instructions. The resulting template was loaded on an Ion 316™ Chip and sequenced on an Ion PGM™ sequencer using an Ion PGM™ 200 Sequencing Kit (Life Technologies, Carlsbad, CA, USA).

### Sanger sequencing

Primers for Sanger sequencing were designed to cover targeted regions of the AmpliSeq™ Panel. Thirty-eight primer pairs covered 11876 bp of AmpliSeq™ Panel, approximately 68% of all NGS targeted regions (**Supplementary Table S1**). Sanger sequencing was performed on an Applied Biosystems ABI3730xl DNA analyzer by the Microsynth laboratory (Balgach, Switzerland). Sanger sequencing traces were aligned to target reference sequences, and variants were detected using Mutation Surveyor (MS) software v.4.0.9 (SoftGenetics, State College, PA, USA) with default parameters for bidirectional sequencing data.

### Samples

Characterized clinical samples with one of the following diagnoses: cystic fibrosis, phenylketonuria and galactosemia, were provided by Federal State Budgetary Institution “Research Centre for Medical Genetics”, Moscow and The Research Institute of Medical Genetics of the Russian Academy of Medical Sciences, Tomsk. Control samples from healthy individuals were collected by Parseq Lab. A total number of unique samples obtained for the clinical trials was 315. Informed consent was obtained for all cases. The Institutional Review Board of RCMG had approved this study.

## RESULTS

### Assay design

We have analyzed open-source locus-specific databases, molecular diagnostics guidelines and scientific publications in order to select clinically significant variants in studied genes: *CFTR*, *PAH* and *GALT*. We have developed a supervised SeqDB database containing more than 600 selected variants with annotations including variant’s trivial name, genetic structure, clinical information and external references (**Supplementary materials S2**). In order to achieve unambiguous variant identification SeqDB operates with variants via developed mutation key (**Fig.1**). SeqDB variants (hotspots) together with functional gene regions were used for the selection of target regions for the panel (**Fig.2**). The final coverage of target regions by AmpliSeq™ designer comprised 99%. Only last six amino acids of exon 21 CFTR carrying two rare CFTR variants 3600G>A (c.3468G>A) and D1152H (c.3454G>C) are not covered by the Panel. These variants are included in the assay limitations list.

**Fig. 1.**
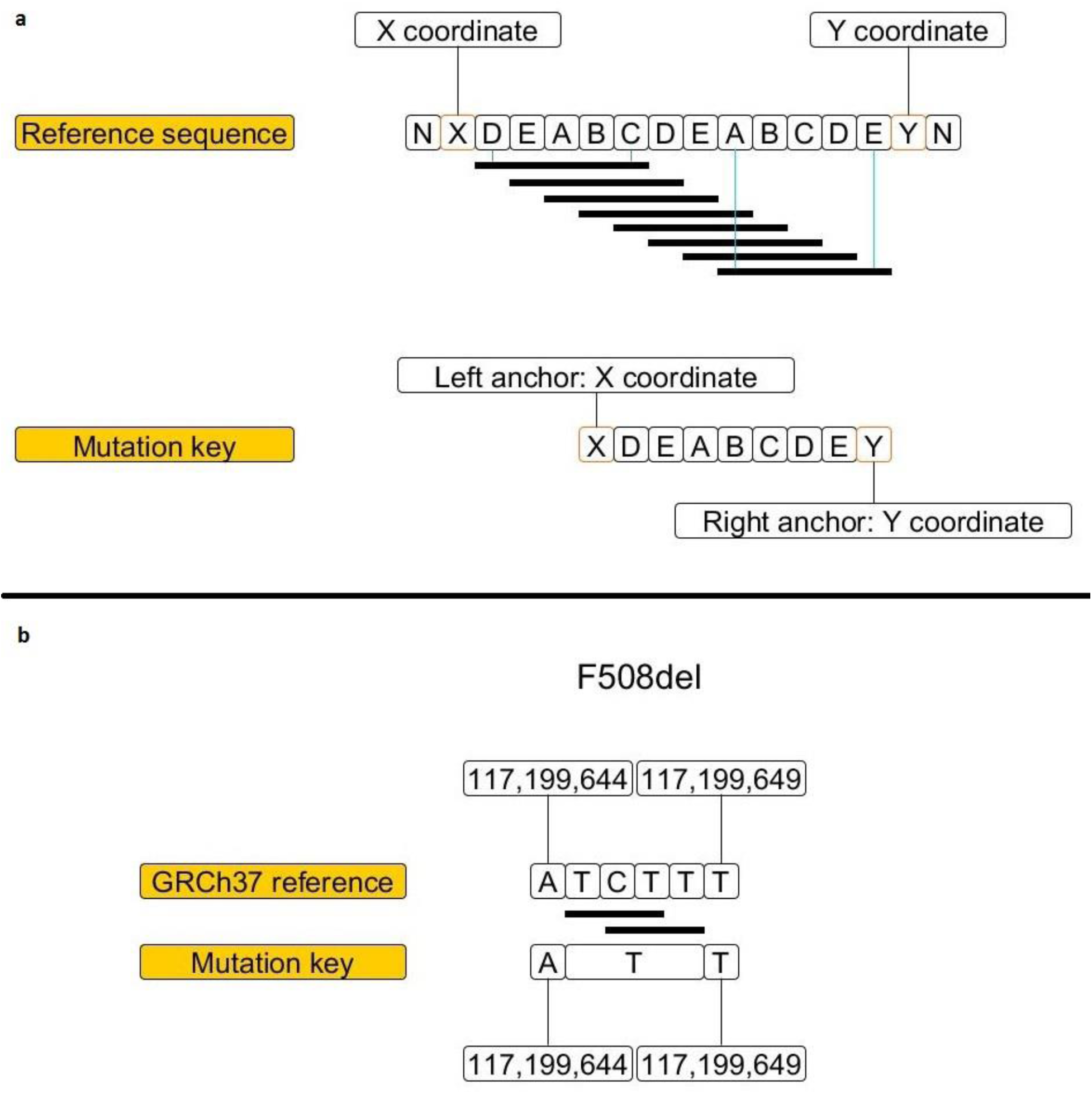
Mutation key. (**a**) Ambiguity of variant identification in repetitive sequences. Deletion of 5bp, DEABC, can be designated as 8 different variants. (**b**) Unambiguous variant identification using mutation key algorithm, on F508del example. Explicitness is achieved via two anchors corresponding to an unaltered bases and the resulted sequence between them. The mutation key is the same for two possible records.

**Fig. 2.**
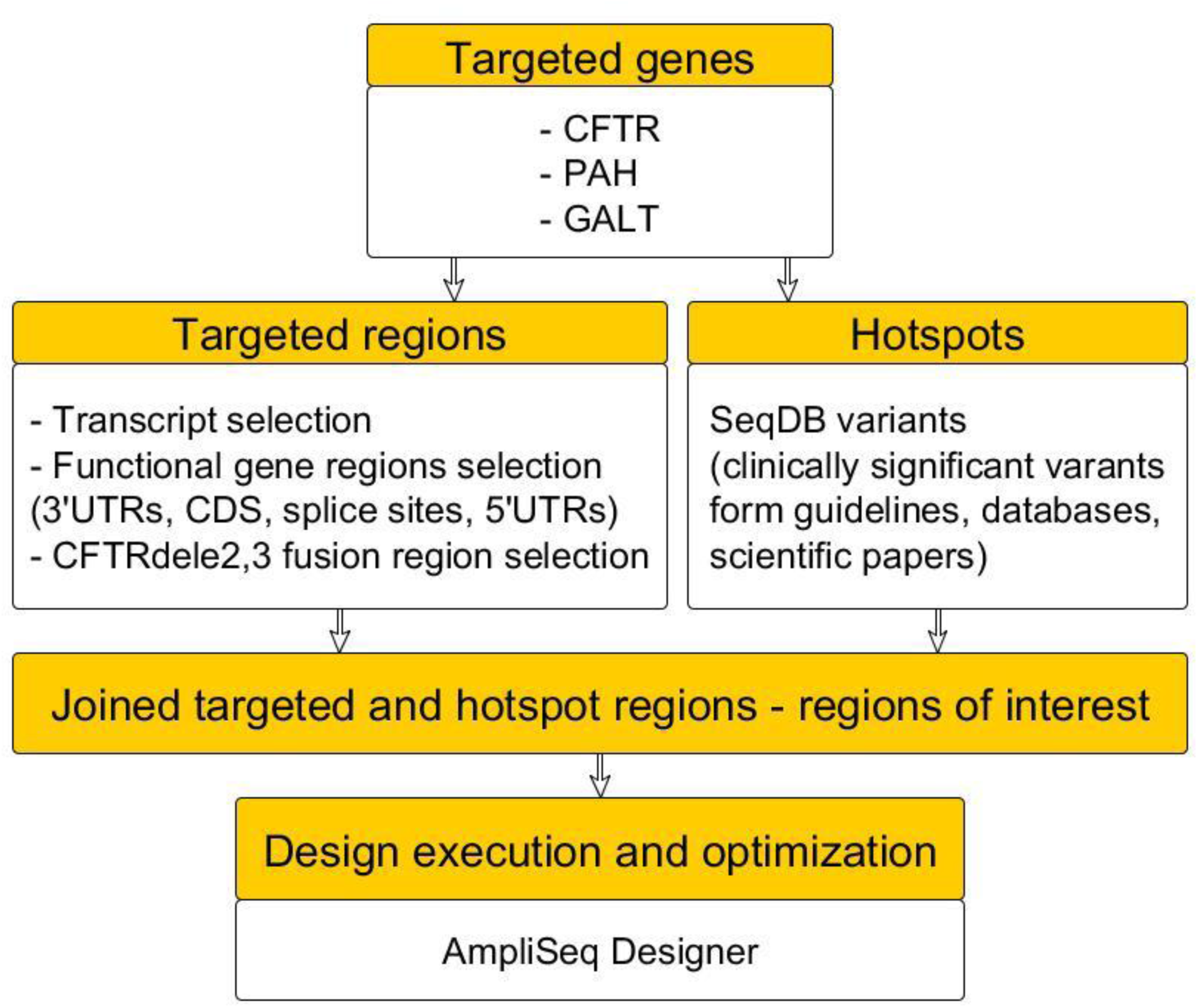
Panel design. Regions of interest were selected by joining functional regions of the genes and known clinically significant variants in order to be sure that all relevant biomarkers are covered by the Panel.

Data analysis comprises two steps. Initial data processing including basecalling, alignment and variant calling is performed on the Torrent Server™ with customized plugins settings. Secondary data analysis including data quality control, variants annotation and report generation is performed using the developed VariFind™ software.

Quality control is based on the determination of sequencing quality across the full length of targeted region. Sequencing quality of the region is measured by the quality corrected coverage (QCC) of the region:

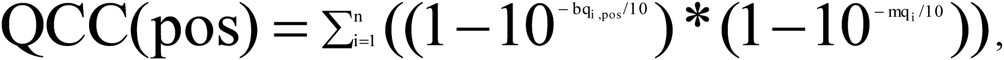

where: *i* - read number, *n* - coverage, *bq* - phred-base quality, *mq* - mapping quality.

Sample’s sequencing data quality cutoff is set by the proportion of targeted regions with QCC above the selected threshold.

Variants annotation consists in identification of clinically significant variants, i.e. variants included in SeqDB™ database. For all clinically significant variants, the detailed annotations are available (**Supplementary materials S3**). For other identified variants, the dbSNP annotation and HGVS nomenclature are provided. Since the sequencing coverage is uneven, that is typical for all sequencing platforms, and some of targeted regions may be low-covered, the software provides information on sequencing quality in the wild type positions corresponding to the clinically significant variants.

VariFind™ software generates the analytical report containing the following information: laboratory’s contact information, patient’ data, sample ID and type, name of the technician conducting the assay, date of sequencing, the aim of testing, testing genes, assay method, assay’s performance characteristics, table of identified clinically significant variants, table of other identified variants, SeqDB database version used for the annotation, proportion of targeted regions covered by the assay and clinically significant variants that are not covered by the assay in the sample, date of report generation and the name of molecular pathologist generated the report (**Supplementary materials S4**). Identified variants are provided with variant name or dbSNP ID, HGVS nomenclature and zygosity. Comments of molecular pathologist are also included in the report if present. The analytical report is then interpreted by the medical geneticist or genetic counsellor who generates the medical conclusion upon the assay results.

### Analytical validation

We developed analytical validation protocol for NGS data based on the recommendations and guidelines for clinical laboratories issued by Swiss Society of Medical Genetic (**SSMG 2003)**, National Genetics Reference Laboratory (**NGRL 2007**), Clinical and laboratory standards institute (**CLSI 2008**), American College of Medical Genetics and Genomics (**ACMG 2013**) and US Centers for Disease Control and Prevention (**CDC 2015**). Analytical validation consisted in pairwise base comparison between the NGS and Sanger sequencing across the full length of targeted NGS region (**Fig.5**). Data comparison and calculation of analytical characteristics were performed using in-house developed script written in Python. All calculations were made considering the genome is diploid, i.e. homozygous variants is considered as two independent variants.

Analytical sensitivity was calculated as the proportion of common variants identified by NGS and Sanger sequencing among all Sanger-specific variants. Analytical specificity was calculated as the proportion of common variants identified by NGS and Sanger among all NGS-specific variants. General accuracy was calculated as the proportion of common base calls between Sanger sequencing and NGS among all base calls within the sequenced region.

Analytical validation was carried out using 89 clinical samples and 10 control samples. Total number of analyzed Sanger sequencing traces is 7178. Comparison is performed by 1466794 positions sequenced by both methods. The number of variants identified is 774, of which 769 is common. All discrepancies were due to the different basecalls identified on the forward and reverse strands by the Mutation Surveyor. Manual analysis confirmed results obtained by the NGS (**Supplementary materials S5**). Nevertheless, according to the validation protocol, we had calculated these differences as false-negatives and false-positives for NGS. Thus, analytical sensitivity, analytical specificity and general accuracy of the assay are comprised 99.87% (95% CI: 99.28-100.00%), 99.23% (95% CI: 98.32-99.72%) and 99.99% (95% CI: 99.99-99.99%), correspondingly.

The accuracy of CNV detection is comprised 100% for CFTRdele2,3 detected by the evaluation of fusion amplicon coverage and 85% for all other CNVs detected using CONVector tool (Parseq Lab).

### Clinical validation

We developed clinical validation protocol for NGS data based on the same recommendations and guidelines as for the analytical validation. Validation consisted in blind testing of clinical and control samples in four independent laboratories. In order to evaluate diagnostics characteristic of the assay clinical samples with two known pathogenic alleles and control samples from healthy individuals were received from biobanks (group 1). In order to evaluate analytical performance CF-clinical samples with only one or without any known pathogenic alleles were received from biobanks (group 2).

Diagnostic sensitivity was calculated as the proportion of the assay positive samples among true positive samples. Diagnostic specificity was calculated as the proportion of the assay negative samples among control samples. Analytical performance was calculated as the proportion of novel pathogenic alleles identified by the assay among all pathogenic alleles.

Total number of group 1 samples tested in reference laboratories is 470 (**Fig.3**). Total number of unique pathogenic mutations tested during validation is comprised 21, 17 and 7 for CF, PKU and GAL correspondingly (**Supplementary materials S6**). During the validation, two false negative and zero false positive results were obtained. Both false negative results aroused from one laboratory and resulted from the cross-contamination of two samples during the library preparation step. Thus, diagnostic sensitivity and specificity comprised 99.36% (95% CI: 97.71-99.92%) and 100.00% (95% CI: 98.72-100.00%), correspondingly.

**Fig. 3.**
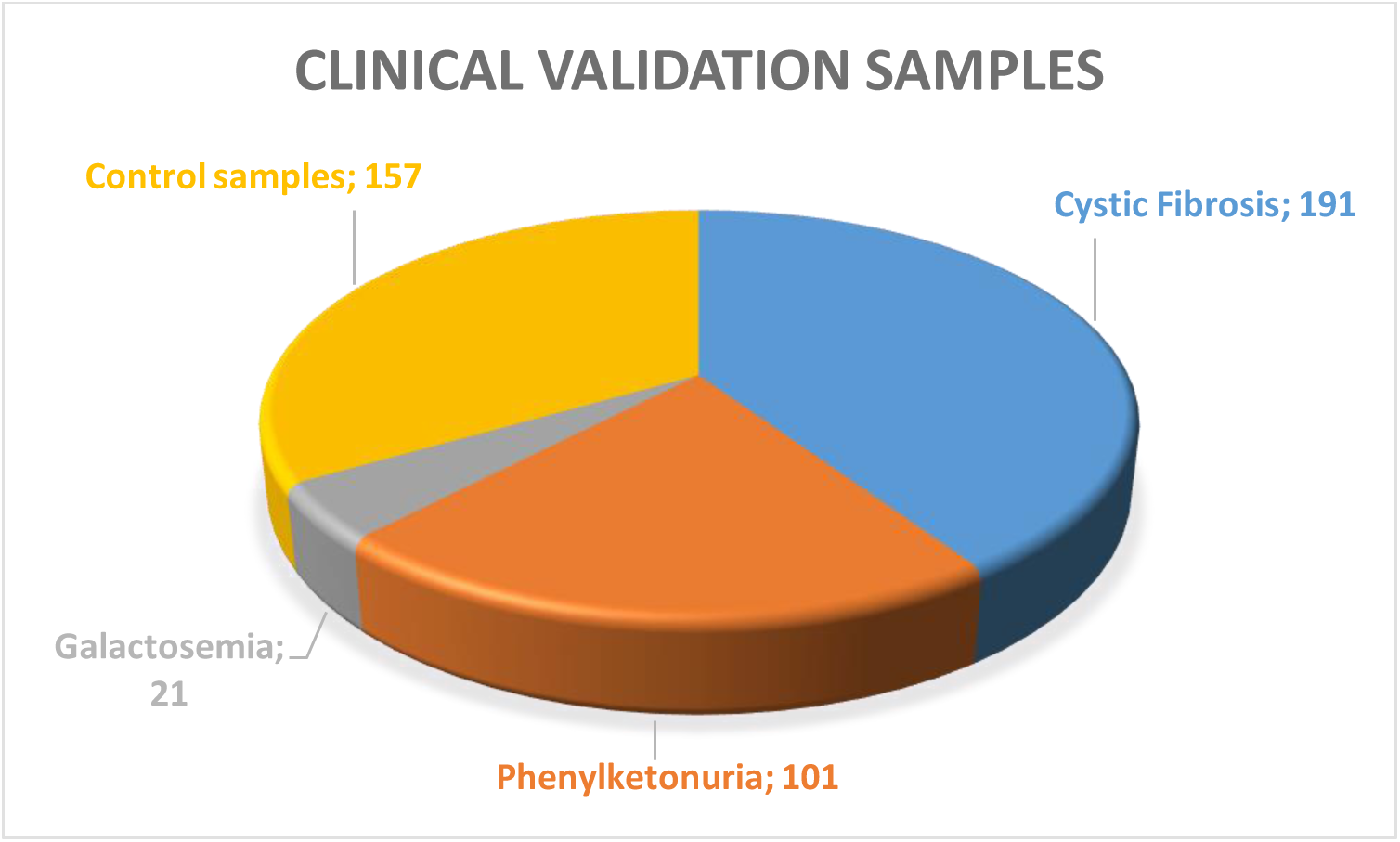
Group 1 samples tested during clinical validation.

Using NGS-based assay 48 a priori known variants have been confirmed and 44 novel variants have been identified (**Supplementary materials S7**). Thus, analytical performance is comprised 47.82%.

Interlaboratory reproducibility was evaluated using 124 samples tested in three different laboratories. Among 372 obtained results, 4128 variants have been identified in three laboratories, 4124 of them were common. Discordant variants were identified in two samples that were cross-contaminated in one reference laboratory. Thus, reproducibility comprised 99.90% (95% CI: 99.75-99.97%).

### Implementation

After the validation has been completed, the assay is being implemented in clinical laboratory for the confirmatory diagnostics. Over the 10 months, 203 samples have been tested. In 81% of CF-suspected samples and 61% of PKU-suspected samples, at least one pathogenic allele has been detected. The inability to detect mutations in all patients can be explained by deep intronic mutations and mutations in modifying genes not included in the panel (**Castellani 2008**). This may be the basis for further panel modifications.

Regardless of such relatively limited clinical usage of the assay ten novel pathogenic alleles have been identified in the *CFTR* gene and one - in the *PAH* gene (**Table 1**). All these variants were described in patients with corresponding clinical diagnosis and were confirmed by Sanger sequencing.

**Table 1.** Novel mutations identified in CF and PKU patients.

| # | Gene | Mutation | HGVS name | Genotype | Diagnosis |
| --- | --- | --- | --- | --- | --- |
| 1 | PAH | 1298dupA | c.1294_1295insT | R158Q/1298dupA | PKU |
| 2 | CFTR | 1812-1G>C | c.1680-1G>C | R334W/1812-1G>C | CF |
| 3 | CFTR | 3272-16T>A | c.3140-16T>A | F508del/3272-16T>A | CF |
| 4 | CFTR | 440del8 | c.308_315delGAAGAA<br>TC | F508del/440del8 | CF |
| 5 | CFTR | 663dupT | c.531_532insT | F508del/663dupT | CF |
| 6 | CFTR | I530T | c.1589T>C | F508del/I530T | CF |
| 7 | CFTR | IVS22+6T>A | c.4136+6T>A | R31C/IVS22+6T>A | CF |
| 8 | CFTR | L581X | c.1742T>G | 406-1G>A/L581X | CF |
| 9 | CFTR | L863R | c.2588T>G | F508del/L863R | CF |
| 10 | CFTR | Q378X | c.1132C>T | F508del/Q378X | CF |
| 11 | CFTR | W361X | c.1083G>A | F508del/W361X | CF |

## DISCUSSION

NGS is a rapidly evolving technology allowing extracting valuable genetic information from vast genomic regions. It is now became affordable for the routine clinical usage. Translational studies are published for different NGS clinical applications and many laboratories offer laboratory-developed tests (LDT). However, the usage of new technology in a routine clinical practice is still limited due to the fundamental differences of NGS technology as compared with classic molecular genetic methods. The size of the genome region tested complicates the assay validation and quality control. Practically unlimited number of variants that can be identified with the assay drastically challenges the results interpretation.

In this work, we attempted to develop a comprehensive ready-to-use NGS-based assay kit. We developed user-friendly software for clinician, not necessary to be familiar with bioinformatics. We developed QC-procedure that allows easy filtering out samples not suitable for the downstream analysis. Supervised SeqDB™ database, that integrates information from different annotation sources, facilitates the interpretation. Another advantage of SeqDB™ is the possibility to extend the number of reportable biomarkers just by adding the variants to the database with the appearance of new scientific data.

We performed validation in accordance with the clinical trials standards and obtained CE-IVD certification for the assay. The assay kit includes reagents, control sample, software and the user manual.

The assay is being implemented in a clinical laboratory during a year and revealed successful performance for confirmatory diagnostics. The assay is especially useful for genetically heterogenic populations, such as multi-ethnic Russian population.

The clinical utility of the assay is also under determining within the framework of the Russian Newborn Screening program in St-Petersburg. Started in 2015 as a pilot clinical approbation, the assay is being used as the second stage of a second-tier newborn screening test to assess the effectiveness of NGS-based assays in facilitation the diagnostics of CF, PKU and GAL in newborns. The results of the upcoming study will be used to evaluate the healthcare, social and economic perspectives of including NGS-based DNA diagnostics into the newborn screening program.

